# The fruitENCODE project sheds light on the genetic and epigenetic basis of convergent evolution of climacteric fruit ripening

**DOI:** 10.1101/231258

**Authors:** Peitao Lü, Sheng Yu, Ning Zhu, Yun-Ru Chen, Biyan Zhou, Yu Pan, David Tzeng, Joao Paulo Fabi, Jason Argyris, Jordi Garcia-Mas, Nengui Ye, Jianhua Zhang, Donald Grierson, Jenny Xiang, Zhangjun Fei, Jim Giovannoni, Silin Zhong

**Affiliations:** State Key Laboratory of Agrobiotechnology, School of Life Sciences, The Chinese University of Hong Kong, Hong Kong, China; College of Horticulture, South China Agricultural University, Guangzhou, China; College of Horticulture and Landscape Architecture, Southwest University, China; Department of Food Science and Experimental Nutrition, FCF, University of Sao Paulo, Sao Paulo, Brazil; IRTA, Centre for Research in Agricultural Genomics, Barcelona, Spain; Department of Biology, Hong Kong Baptist University. Hong Kong, China; School of Crop Sciences, University of Nottingham, UK, Zhejiang Provincial Key Laboratory of Horticultural Plant Integrative Biology, Zhejiang University, China; Weill Medical College, Cornell University, New York, NY 10021, USA; Boyce Thompson Institute for Plant Research, Cornell University, New York, USA

## Abstract

Fleshy fruit evolved independently multiple times during angiosperm history. Many climacteric fruits utilize the hormone ethylene to regulate ripening. The fruitENCODE project shows there are multiple evolutionary origins of the regulatory circuits that govern climacteric fruit ripening. Eudicot climacteric fruits with recent whole-genome duplications (WGDs) evolved their ripening regulatory systems using the duplicated floral identity genes, while others without WGD utilised carpel senescence genes. The monocot banana uses both leaf senescence and duplicated floral-identity genes, forming two interconnected regulatory circuits. H3K27me3 plays a conserved role in restricting the expression of key ripening regulators and their direct orthologs in both the ancestral dry fruit and non-climacteric fleshy fruit species. Our findings suggest that evolution of climacteric ripening was constrained by limited availability of signalling molecules and genetic and epigenetic materials, and WGD provided new resources for plants to circumvent this limit. Understanding these different ripening mechanisms makes it possible to design tailor-made ripening traits to improve quality, yield and minimize postharvest losses.

**One Sentence Summary:** The fruitENCODE project discovered three evolutionary origins of the regulatory circuits that govern climacteric fruit ripening.

## Introduction

Fleshy fruit ripening is a developmental process unique to flowering plants, in which the chemical and physiological properties such as colour, aroma, flavour, texture and nutritional contents of the seed-bearing organ are dramatically altered, in order to attract frugivores as seed dispersers (*1, 2*). As vital to human and animal diets, understanding fruit ripening is increasingly important for global food and nutritional security.

Many economically important fruits such as apple, banana and tomato can be harvested mature but unripe, stored and treated with the plant hormone ethylene to complete the ripening process. They show a concomitant increase in respiration and ethylene biosynthesis upon initiation of ripening, characteristic of so-called climacteric fruits. Non-climacteric fruits like grape and strawberry do not display these features and has a less clear requirement of ethylene for ripening.

Fleshy fruit is a classic example of convergent evolution, and it has occurred multiple times in the history of angiosperms. However, its molecular basis remained largely unknown, as most of the studies have focused on a single model fruit tomato (*3*). Tomato produces ethylene at the onset of ripening, and disrupting ethylene biosynthesis or perception inhibits ripening (*4*). Besides ethylene, a series of transcription factors such as COLORLESS NONRIPENING (CNR), NON-RIPENING (NOR) and RIPENING INHIBITOR (RIN) are also required for tomato fruit ripening (*3, 5, 6*). Among them, the MADS-box transcription factor RIN is most extensive studied, and its mutant alleles are widely used in commercial tomato production. The MADS gene family is present in all eukaryotes and has expanded considerably in plants. *RIN* belongs to the MIKC-clade, members of which are involved in floral organ development. RIN physically interacts with additional MADS-box proteins and can bind directly to the promoter of key ripening genes, including those involved in colour change, fruit softening, aroma volatile production and sugar metabolism, and is also responsible for regulating ethylene biosynthesis genes such as *ACO1* and *ACS2*, which are induced during fruit ripening and encode the committed steps of ethylene synthesis (*7*).

In ripening tomato fruit, ethylene biosynthesis is autocatalytic, which is historically referred to as system II ethylene, to be distinguished from the self-inhibitory system I ethylene production in other tissues including immature fruit (*8*). The autocatalytic nature suggests that there is a positive feedback loop controlling ethylene production during fruit ripening.

However, ethylene is a stress hormone involved in many plant developmental processes including, for example, tissue senescence and abscission (*4*). Such an autocatalytic system involving a diffusible signal molecule poses a major threat to the plant itself and its neighbours, as any leakiness could cause developmental perturbtions or even tissue death. It has been shown that dynamic DNA methylation changes plays an important role in restricting the expression of tomato ripening genes (*5, 7*). Many ripening gene promoters bound by RIN are demethylated during fruit development, and silencing the fruit-specific DNA demethylase *DML2* can inhibit ripening (*9*).

While tomato is still the predominant model for ripening research, many fleshy fruit genomes have now been sequenced. Tomato is also the only species reported so far to induce a DNA demethylase expression during fruit ripening (*9*), begging the question to what degree that the tomato model is universal. In addition, tomato has experienced whole-genome duplication (WGD) ~71 Myr, and key ripening regulators including *RIN* and ethylene biosynthesis genes *ACO1* and *ACS2* are paralog members of duplicated gene families (*10*). Hence, other species without a ripening-specific demethylase and WGD might have evolved different regulatory approaches to facilitate and coordinate the ethylene-dependent ripening phenomena.

Some fruits also have unique ripening traits that are absent in tomato. For example, both climacteric and non-climacteric genotypes exist in melon and pear (*11, 12*). It has also been suggested that banana operates an additional leaf-like self-inhibitory ethylene system I during ripening, and unlike tomato, once its autocatalytic ethylene production starts, it becomes ethylene-independent (*13*).

It is not possible to resolve complex convergent traits in diverse taxa such as fleshy fruit ripening by sequencing and comparing genomes if the convergence occurred through the evolution of different genes and pathways, or if the regulatory interactions are different. To tackle this, we have used an ENCODE style functional genomic approach to systematically characterize the molecular circuits controlling ripening in 13 plant species. We found three major circuits adopted by different plants in the evolution of climacteric fruit. Part of the genes and epigenetic marks are surprisingly conserved even in dry and non-climacteric species, suggesting that these different ethylene-dependent fleshy fruit ripening mechanisms are evolved from existing pathways in the ancestral angiosperm’s leaf and carpel tissues.

## Results

### The fruitENCODE data

Our current fruitENCODE dataset comprises seven climacteric fruit species (apple, banana, melon, papaya, peach, pear and tomato), four non-climacteric species (cucumber, grape, strawberry and watermelon) and two dry fruit species *(Arabidopsis* and rice) (tables S1 to S18). We have generated 361 transcriptome profiles from their leaves and fruit tissues at different developmental stages. We have also generated 66 tissue-specific accessibility chromatin profiles, and identified between 36,584 to 45,116 DHSs that could be associated to 43.28% to 71.93% of annotated genes in each species. The cis-regulatory elements in the DHSs enabled us to predict and experimentally validate the regulatory circuits controlling climacteric fruit ripening, and further recreate each circuit in tobacco leave. In addition, we have generated 43 methylomes and 76 histone modification profiles to investigate which epigenetic factors are responsible for regulating these ripening circuits. All datasets can be accessed from the fruitENCODE database.

### Tomato, apple and pear fruits use a MADS positive feedback loop

We first reconstructed the ripening regulatory circuit in the model fruit tomato that has three key components: ethylene, transcription factor RIN and DNA methylation. The main molecular features of ethylene signalling are conserved in plants and its signal perception by the receptors leads to the stabilization of the EIN3-type transcription factors, which activate the downstream genes (*4*). From the open chromatin dataset, we found an EIN3 binding motif in the promoter of *RIN*, the functional significance of which was confirmed by our EIN3 ChIP-Seq (**Fig. 1A**). RIN is a MADS-box transcription factor, which functions in a multimeric complex with TAGL1, which is constantly expressed during both early and late fruit development *(14)*. We have also performed ChIP-Seq for RIN and TAGL1, and found that they can directly target the rate-limiting ripening ethylene synthesis genes *ACS2* and *ACO1* (**Fig. 1A**). In addition, our DHS and methylome data showed that the EIN3 binding site in the *RIN* promoter is demethylated and becomes accessible in ripening fruit tissues (**Fig. 1A**).

**Fig. 1.**
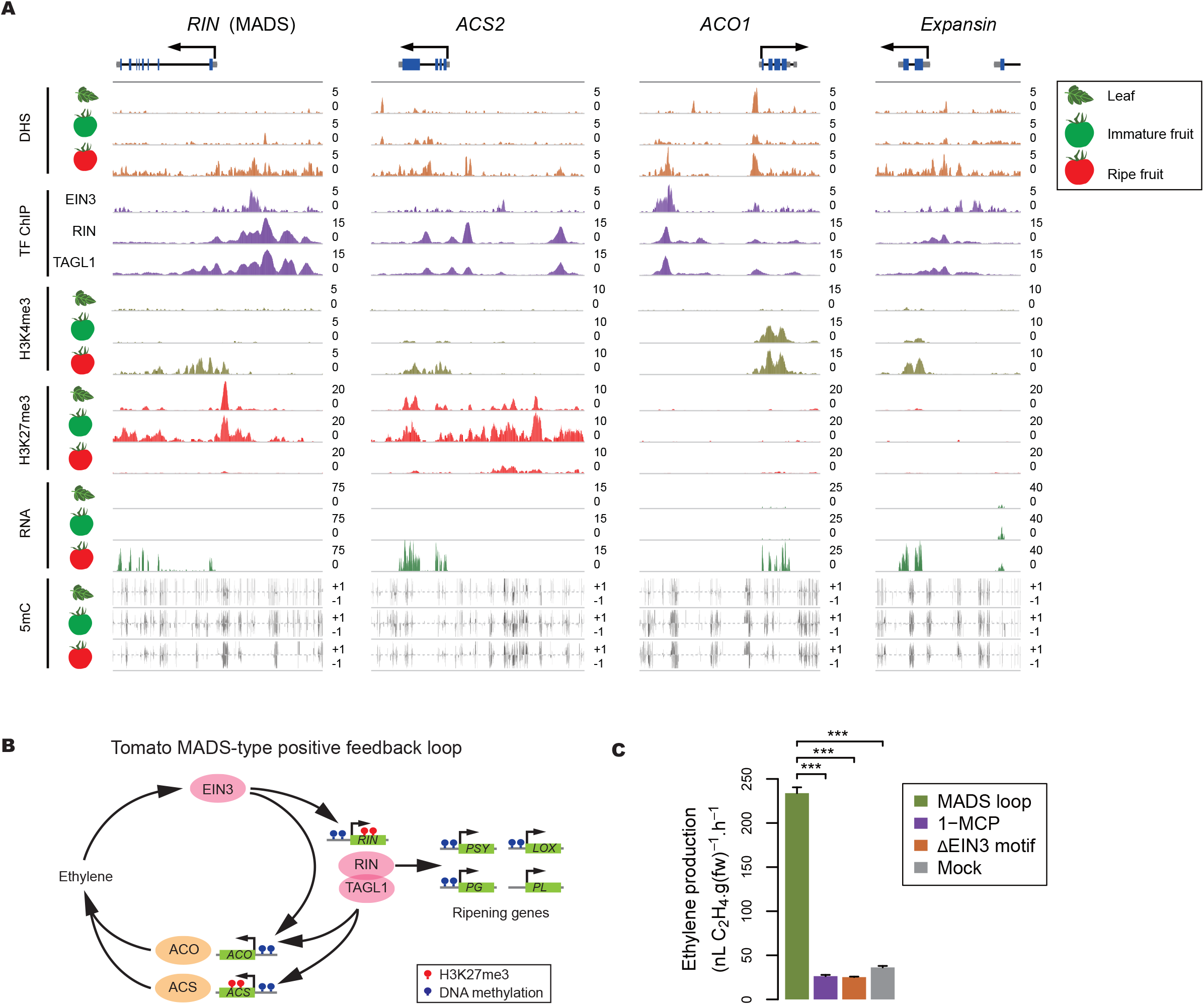
MADS positive feedback loop controlling tomato fruit ripening. (**A**) Gene loci involved in fruit ripening with browser tracks showing different epigenetic features, transcription factor binding and gene expression. (**B**) Model for tomato fruit ripening regulation. In nonfruit tissues and immature fruits, key ripening regulators are repressed by promoter DNA hypermethylation and H3K27me3. When the epigenetic marks are removed in the matured fruits, ripening is initiated with a burst of autocatalytic ethylene production, followed by the activation of the downstream ripening genes. (**C**) Ectopic expression of *RIN, TAGL1* and *ACS2* generated autocatalytic ethylene in tobacco. Ethylene inhibitor treatment or deleting the EIN3 binding motif in the *RIN* promoter disrupted the loop and blocked ethylene synthesis.

Our findings suggest that EIN3 and RIN-TAGL1 form a positive feedback loop to synthesize the autocatalytic system II ethylene during tomato fruit ripening (**Fig. 1B**). In addition, ripening genes are also coupled to the loop through RIN-TAGL1. Our ChIP-Seq data showed that the RIN-TAGL1 complex targets well-known ripening genes such as *EXPANSIN, PG* and *PL* that are involved in fruit softening, *PSY, LOX* and *SuSy* that are involved in colour change, aroma production and sugar metabolism, respectively (**table S28 and section 10.5**). Given the central role of the MADS-box transcription factors in this ripening model, we named it the MADS positive feedback loop.

To test this MADS positive feedback loop, we used the tobacco leaf transient expression system that has a functional ethylene signalling pathway and endogenous EIN3 and ACO activities, but lacks the RIN-TAGL1 and ACS activity. When we expressed the tomato *ACS2* and *RIN* under their native promoters, and *TAGL1* was supplied under a constitutive CaMV35S promoter, spontaneous ethylene synthesis was observed (**Fig. 1C**). Addition of 1-MCP, a competitive inhibitor of ethylene, or deletion of the EIN3 binding motif in the *RIN* promoter, interrupted the positive feedback loop and prevented ethylene synthesis, confirming the autocatalytic nature of the regulation of ethylene synthesis (**Fig. 1D**).

We have examined transcriptome and open chromatin data from other species and found that apple and pear have similar ripening-fruit-specific MADS genes with EIN3 binding motifs, while their ethylene biosynthesis genes *ACS* and *ACO* have MADS transcription factor binding motifs. These observations suggests that they could operate a tomato-like MADS positive feedback loop (**figs. S1 and S2**). We have used the tobacco leaf system to test whether the apple and pear genes could generate autocatalytic ethylene. The results showed that their MADS loops are indeed functional, and EIN3 motif deletion confirmed that the EIN3 and these MADS genes could formed a positive feedback loop (**figs. S1 and S2**).

### Key ripening genes are associated with conserved Polycomb H3K27me3 marks

Since ethylene is a stress hormone, it is vital for the plant to repress this loop in non-ripening tissues and confine autocatalytic ethylene action to specific tissues or organs, such as ripening fruits, abscission zones and senescing leaves. It has been shown that whole-genome demethylation is required for tomato fruit ripening (*7, 9*). However, our RNA-Seq data showed that the expression of the apple and pear demethylases is not ripening-specific, neither are the demethylase of other fleshy fruits species we examined (**table S16**).

Our methylome datasets confirmed that there is no tomato-like global demethylation event in apple and pear (fig. S6). Although we could identify hundreds of ripening genes with promoter DMRs in all seven climacteric fruits we studied (**tables S10 to S12**), only in tomato is the master ripening regulator in the positive feedback loop, MADS-box *RIN* associated with DMR. For example, we have identified 11,911, 17,231 and 17,455 genes associated with promoter hypo-DMR in tomato, apple and pear, respectively (**table S12**). In apple and tomato, both *PSY* and *PG*, which are involved in colour change and fruit softening, are associated with promoter hypo-DMR. However, unlike tomato, the two apple MADS-box transcription factor genes in the positive feedback loops are associated with promoter DMR (**fig. S1**). These results suggest that although DNA methylation change in the promoter DHS is widespread, it might not be a conserved mechanism utilized by climacteric fruits to regulate ripening genes.

However, from the histone modification datasets, we found that tomato, apple and pear have the repressive histone mark H3K27me3 on their ethylene biosynthesis and MADS gene loci, whereas they are removed in the ripe fruit tissues (**Fig. 1A and figs. S1 and S2**). The fruitENCODE data also includes multiple cultivars and mutants, and we found that the ethylene-independent pear cultivar Dangshanshuli contains hyper-H3K27me3 in its *ACS* locus compared to the ethylene-dependent cultivar Williams (**fig. S2**). From the tomato ripening mutant collection, we also observed that the non-ripening mutant *nor* contains hyper-H3K27me3 in the *ACS2* and *RIN* loci (**fig. S23**). These suggests that H3K27me3, instead of DNA methylation, might play a conserved role in regulating the MADS type ripening positive feedback loop.

### Peach, papaya and melon operate a NAC positive feedback loop

Tomato has a WGD ~71 Mya, while apple and pear have a common WGD ~40 Mya. These plants have a similar positive feedback loop using paralog member of the duplicated SEP-clade MADS transcription factor family that controls floral organ development, suggesting that neofunctionalization following WGD could have played a vital part in the evolution of MADS-type climacteric fruit ripening (**fig. S3**). However, climacteric fruits such as peach, papaya and some climacteric melon cultivars can also produce and require autocatalytic ethylene for ripening, and did not undergo recent WGD. Their MADS gene direct orthologs of the tomato ripening regulators, are constantly expressed during fruit development (**tables 23 to 25**), suggesting that if they have evolved a positive feedback loop to synthesize the ripening ethylene, it will require different transcription factors.

We have examined the expression pattern of the papaya, peach and melon ethylene biosynthesis genes, and identified their *ACS* and *ACO* genes that are specifically expressed during fruit ripening (**figs. S24 and S25, table S17**). Also from the DHS datasets, we found NAC transcription factor binding motifs in those *ACS* and *ACO* gene promoters. Intriguingly, all three species have a *NAC* gene with ripening-specific expression pattern similar to the *MADS-RIN* genes in tomato (**fig. S4** and **tables S23 to 25**). NAC is one of the largest plant-specific transcription factor families with members involved in many developmental processes such as senescence, stress, cell wall formation, and embryo development. All three ripening *NAC* genes we identified are direct orthologs of the *Arabidopsis* carpel senescence-related *NAC* transcription factors *NARS1/2*, and are distantly related to the leaf-senescence-related *AtNAP (15, 16)* (**fig. S4**). In addition, we have found EIN3 motif in the promoter DHSs of these *NAC*, suggesting that instead of relying on the duplicated floral identity *MADS* gene, plants without WGD might have evolved a positive feedback loop using the existing carpel senescence *NAC* genes (**Fig. 2**).

**Fig. 2.**
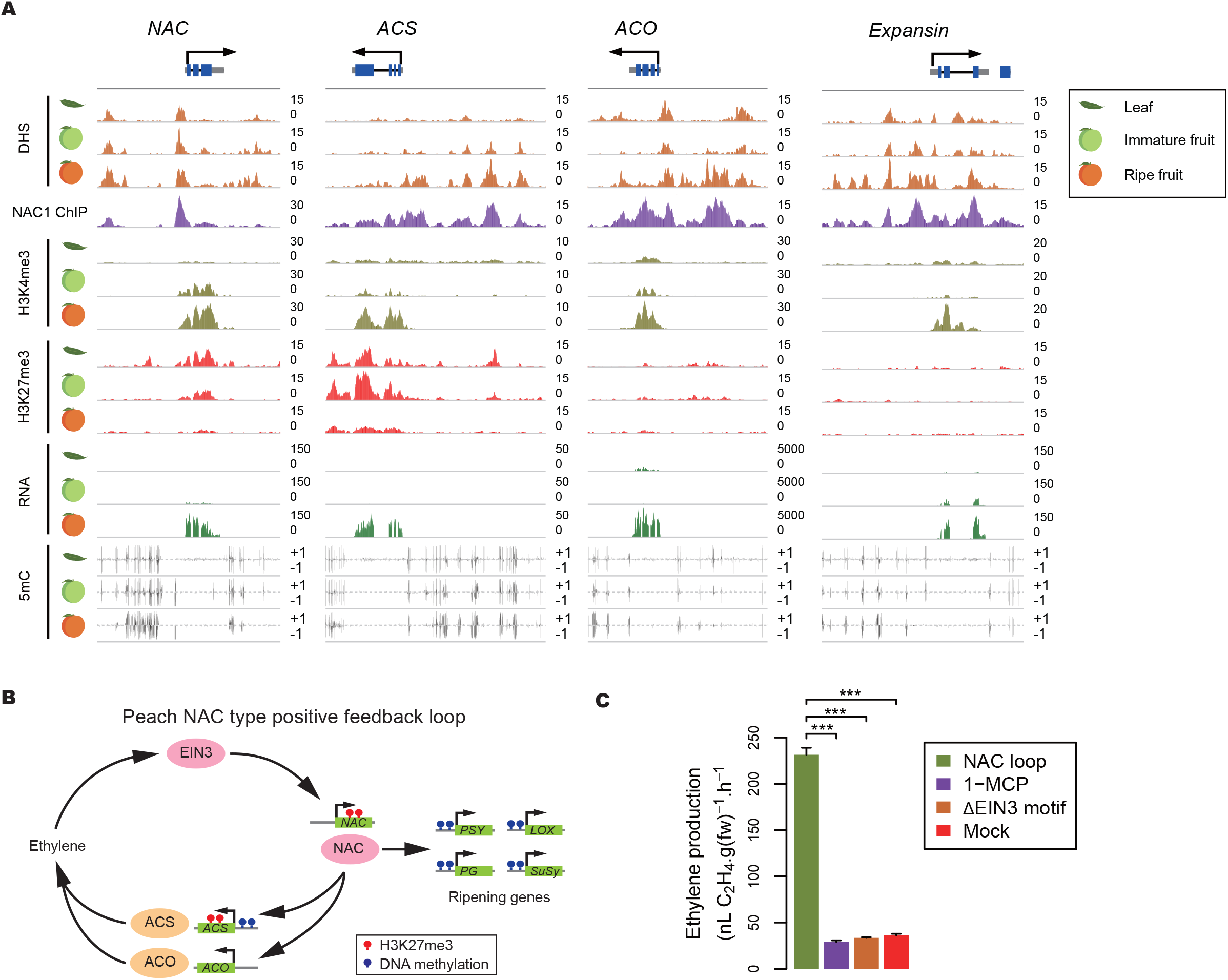
NAC positive feedback loop controlling peach fruit ripening. (**A**) Key gene loci involved in peach ripening with browser tracks showing different epigenetic features, transcription factor binding and gene expression. (**B**), Model for peach fruit ripening regulation. Similar to the tomato model, the peach ripening genes are also repressed by promoter DNA hypermethylation and H3K27me3. (**C**), Ectopic expression of the peach *NAC* and *ACS* genes generated autocatalytic ethylene in tobacco. Ethylene inhibitor treatment or deleting the EIN3 binding motif in the *NAC* promoter disrupted the loop and blocked ethylene synthesis.

To test this, we have ectopically expressed the peach *NAC* (ppa007577m) and *ACS* (ppa004774m) genes under their native promoters in tobacco leaves, and found that they are capable of generating ethylene (**Fig. 2B, C**). The ethylene production could be blocked by treatment with inhibitor 1-MCP, suggesting that it is autocatalytic. We have also used motif deletion assay to confirm that EIN3 binding to the *NAC* gene promoter is required for the autocatalytic ethylene production. These suggest that peach operates a NAC-type positive feedback loop.

In tomato, the downstream ripening genes are directly coupled to the positive feedback loop through the MADS-box transcription factors RIN-TAGL1. To test whether it is the same for plants operating this NAC-type loop, we use ChIP-Seq to identify the direct targets of the peach NAC (**table S25**). Our data showed that it indeed binds to the promoters of the ethylene biosynthesis genes *ACS* and ACO, as well as key fruit ripening genes such as those involved in carotenoid pigment accumulation, volatile secondary metabolite production, cell wall softening and sugar accumulation (**Fig. 2, section 7.5**). The peach DNA methylation and histone modification data showed that genes involved in the NAC positive feedback loop are also associated with the repressive histone mark H3K27me3 in leaves and immature fruit tissues, but not in the ripe fruit (**Fig. 2A**).

In papaya and climacteric melon cultivar Védrantais, we found the same NAC-type positive feedback loops, with key genes associated with H3K27me3 in non-ripening tissues (**figs. S5 and S6**). We also tested their loops using the tobacco system, and performed ethylene inhibitor treatment and EIN3 motif deletion to confirm that they are capable of generating autocatalytic ethylene (**figs. S5 and S6**).

Interestingly, our melon dataset includes the non-climacteric cultivar *Piel de Sapo*, which does not require ethylene for ripening. The QTL *(ETHQV6.3)* responsible for the ethylene-independent phenotype has been mapped to a region, where its *NAC* gene *(MELO3C016540)* involved in the positive feedback loop is located (*17*). It has been shown that non-synonymous mutations in its conserved NAC domain are responsible for a significant delay in the onset of fruit ripening and in the biosynthesis of ethylene in the highly climacteric line Charentais Mono. This NAC gene is also down-regulated in *Piel de Sapo* ripe fruit, when compared to other cultivars, while its promoter is hypermethylated and the genebody is associated hyper-H3K27me3 (**fig. S6**). Similarly, the peach NAC gene involved in the loop is also located in a previously identified QTL associated with the late-ripening phenotype (*18*), suggesting that ripening traits resulting from mutations in the NAC-type loop might have been selected during the breeding process, similar to the widespread used of MADS gene *RIN* mutant allele in the modern tomato cultivars.

### Monocot banana operates a dual-loop system

Banana is also a climacteric fruit that requires autocatalytic ethylene to ripen, and has experienced three recent WGD (*19*). The autocatalytic ethylene production in other climacteric fruits such as tomato and peach can be interrupted by the ethylene-inhibitor, a scenario that we could reproduce in tobacco by ectopic expression of their MADS or NAC-type positive feedback loops. However, inhibitor treatment is unable to interrupt the ethylene production in banana after ripening has been initiated, indicating a transition from autocatalytic to ethylene independent ripening (*13*). Our data suggests that banana has two positive feedback loops, and the second one is able to maintain the ethylene synthesis when the first ethylene-dependent loop is blocked (**Fig. 3**).

**Fig. 3.**
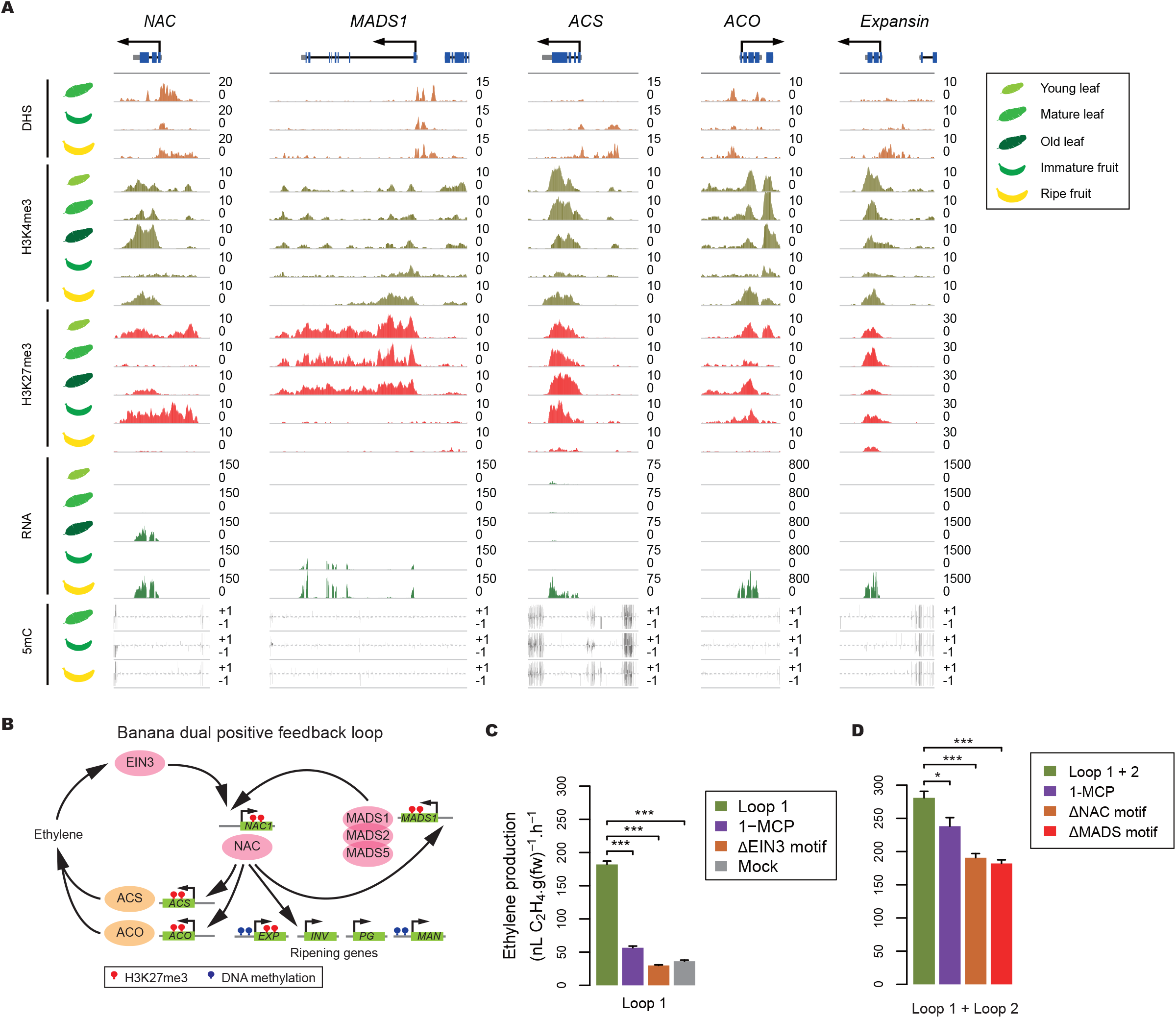
Banana fruit ripening controlled by a dual-loop system. Different from other climacteric fruits, once the banana autocatalytic ethylene production starts, it becomes ethylene independent, and we found that banana has two positive feedback loops to control ripening. (**A**), Genes involved in banana ripening with browser tracks showing different epigenetic features and gene expression. (**B**), Dual-loop model for banana fruit ripening regulation. (**C**), Ectopic expression of the *NAC* and *ACS* genes from the first loop generated autocatalytic ethylene in tobacco. Ethylene inhibitor treatment or deleting the EIN3 binding motif in the *NAC* promoter disrupted the loop and blocked ethylene synthesis. (**D**), Ectopic expression of the *NAC, ACS*, and the three *MADS* genes generated autocatalytic ethylene that could not be blocked by ethylene inhibitor treatment. Deletion of the NAC motif in *MADS1* promoter or the MADS motif in the NAC gene promoter disrupted the second loop.

The first banana loop is similar to the NAC-type positive feedback loop in eudicots such as peach, papaya and melon. We found that the banana NAC transcription factor (Ma06_t33980.1) could activate the ripening ethylene biosynthesis genes *ACS* (Ma04_t35640.1) and *ACO* (Ma07_t19730.1), while its own promoter contains an EIN3 binding motif like peach NAC gene (**Fig. 3B and fig. S22**). To test the first loop, we ectopically expressed the banana *NAC* and *ACS* genes under their native promoters in tobacco and found that they are sufficient to generate ethylene in an autocatalytic manner (**Fig. 3C**). Ethylene inhibitor 1-MCP, as well as EIN3 motif deletion, could block ethylene production in the absence of the second loop, suggesting that the banana loop I is a functional NAC-type positive feedback loop.

It should be noted that this banana ripening *NAC* gene is a direct ortholog of the rice leaf senescence transcription factor OsNAP (*20*), and is distinct from the carpel senescence-related *NAC* genes utilized by the eudicots climacteric fruits mentioned above (**fig. S4**). We have examined the gene expression and histone modifications in young, matured and senescence banana leaves, and found that this *NAC* gene is also expressed during leaf senescence (**Fig. 3A and table S20**). Our epigenome data showed that genes in the banana loop I are associated with similar tissue-specific H3K27me3 marks as those in the eudicots NAC-type climacteric fruits, except that the banana *NAC* gene is not associated with H3K27me3 in some leave developmental stages (**Fig. 3A**).

Both the eudicot MADS- and NAC-type positive feedback loops are directly coupled to the downstream ripening genes, which we have confirmed by ChIP-Seq using tomato and peach as example. Without a suitable banana NAC antibody, we have used the dual luciferase assay in tobacco to confirm that the banana NAC in the positive feedback loop is capable of activating downstream ripening genes such as *EXPANSIN, PG, MAN* and *INV* (**fig. S22**).

Interestingly, utilizing a leaf-instead of a carpel-senescence NAC in banana fruit ripening poses a dilemma for the plant as some ripening genes, such as *EXPANSIN*, are not expressed in leaf. We found that the *EXPANSIN* locus is associated with H3K27me3 marks in leaf, suggesting that banana might rely on epigenetic factors to restrict its expression to ripening fruit tissues (**Fig 3A**). In contrast, we did not observe H3K27me3 targeting the ripening *EXPANSIN* gene in MADS or NAC-type eudicot fruits, as their *MADS* and *NAC* genes are ripening-specific and are not expressed in leaf.

It has been previously shown that three MADS transcription factors (MADS 1/2/5) are involved in banana fruit ripening (*21, 22*). *MADS1/2* and tomato *RIN* are in the SEP-clade, while *MADS5* and the tomato *TAGL1* are in the AG-clade. It is important to note, however, that *MADS1/2* and *MADS5* are not direct orthologs of the tomato *RIN* and *TAGL1* (**fig. S3**). The expression *MADS1/2/5* genes could be detected in both immature and ripening fruits, and they are only associated with H3K27me3 in the leaf tissue (**Fig. 3A**). We found a NAC motif in the banana *MADS1* gene promoter and a MADS motif in the *NAC* promoter, suggesting that they could form a second positive feedback loop to bypass the first loop that is ethylene dependent (**Fig. 3B**).

To test the second loop, we first used the tobacco system to show that the first NAC loop could be blocked by either inhibitor 1-MCP treatment or EIN3 motif deletion (**Fig. 3C**). Adding a second loop with *MADS1* driven by its native promoter and *MADS2/5* with the constitutive 35S promoter enabled the tobacco leaves to synthesize 54.27% more ethylene (p=2.91x10^−5^) (**Fig. 3D**). Most importantly, ethylene inhibitor 1-MCP was unable to block the ethylene production when the second loop was present, mimicking the behaviour of the ripening banana fruit. We have also confirmed that the second loop is dependent on the interaction of the *NAC* and *MADS1* genes by deleting the NAC motif in the promoter of *MADS1*, as well as deleting the MADS motif in the *NAC* promoter (**Fig. 3D**). Taken together, our results showed that banana fruit ripening is controlled by a dual positive feedback loop that consists of both *NAC* and *MADS* genes. These banana *NAC* and *MADS* genes are also associated with tissue-specific H3K27me3 marks. But they are of a different evolutionary origin compared to the NAC and MADS genes used by other eudicot climacteric fruits.

### Direct orthologs of the ripening genes in non-climacteric and dry fruit species

To investigate the possible evolutionary origin of these three types of ripening circuits, we have examined four non-climacteric fruits (cucumber, grape, strawberry and watermelon). None of them have recent WGD. These species have direct orthologs of the carpel senescence *NAC* with H3K27me3 modification change and a ripening-specific gene expression pattern similar to those in the NAC-type climacteric fruit peach, papaya and melon (**Fig. 4A and figs. S7 to S10**). However, their *NAC* genes either lack the EIN3 motif in their promoter DHS, or the NAC motif is absent from their ethylene biosynthesis genes, both factors which would preclude participation in a positive ethylene feedback loop. We have also examined their *MADS* genes that are direct orthologs of the tomato *RIN*, and found that they are expressed in fruits, and are associated with H3K27me3 marks in leaves.

**Fig. 4.**
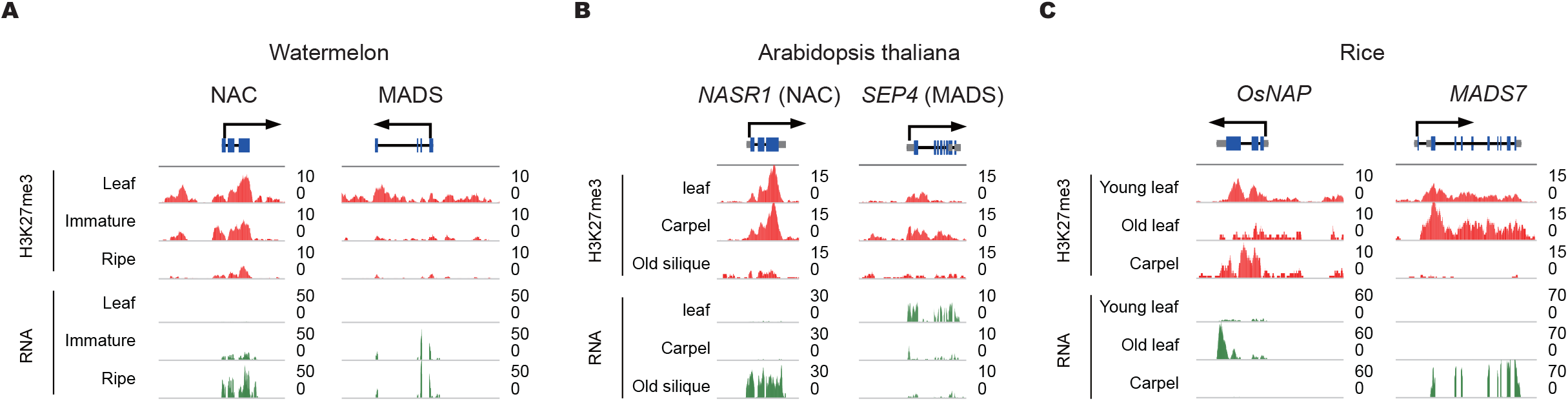
Direct orthologs of the ripening genes in non-climacteric and dry fruit species. The direct orthologs of the eudicot climacteric fruit ripening *NAC* and *MADS* genes in the non-climacteric fruit watermelon (**A**) and dry fruit *Arabidopsis* (**B**) are also associated with tissue-specific H3K27me3 mark. (**C**) The rice direct orthologs of the banana fruit ripening *NAC* and *MADS* genes.

Strawberry appears to be an exception. It possesses all the required cis-regulatory elements in its *NAC (mrna31150.1)* and ethylene biosynthesis gene *ACS (mrna11391.1)* to form a positive feedback loop (**fig. S26**). Its ethylene biosynthesis gene *ACS*, however, is constantly repressed by H3K27me3 during fruit development and ripening (**fig. S9**), again hinting the important roles of the cis-regulatory elements and epigenetic marks in evolving climacteric fruit ripening.

The dry fruit-bearing plant *Arabidopsis* also has direct orthologs of the *NAC* and *MADS* genes that are involved in climacteric fruit ripening. We have examined their H3K27me3 levels in *Arabidopsis* leaves, carpel and senescence silique, which is the equivalent tissue of a ripening fruit (**Fig. 4B**). We found its *NAC* gene (AT3G15510.1) showed reduced H3K27me3 level during silique senescence, while the MADS gene *SEP4* (AT2G03710.1) was also associated with tissue-specific H3K27me3.

The unique dual-loop system used by banana utilizes a direct ortholog of the rice leaf-senescence *OsNAP* gene. We have profiled the gene expression and histone modification in young and senescence rice leaves, as well as its carpel tissues. Our data confirmed that the *OsNAP* gene is expressed in the senescing leaves, and is associated with H3K27me3 in the young leaves and carpel, but not in the senescence leaves (**Fig. 4C**).

Taken together, our finding showed that other eudicot and monocot plants have direct orthologs of the climacteric fruit ripening genes. These genes are involved in leaf senescence, carpel senescence or floral development, and are associated with tissue-specific H3K27me3 marks. As fleshy fruits are believed to have evolved independently in multiple angiosperm lineages, this could suggest that climacteric fruit ripening could have evolved from existing mechanisms controlling leaf and floral tissue gene expression.

## Discussion

The data from the fruitENCODE project shows that there are three major evolutionary paths of the regulatory circuits that govern climacteric fruit ripening (**Fig. 5**). It is common that different species often evolve similar features when exposed to the same selection pressure. However, the probability of complex traits such as climacteric fruit ripening originating multiple times through similar trajectories should be very small, unless there is strong constraint. Hence, it might not be a coincidence that the four eudicot climacteric fruits without recent WGD evolved a NAC-type ripening circuit that can be traced back to carpel senescence, while those with WGD could utilize the duplicated floral identity genes. We hypothesize that using the existing carpel senescence pathway, acquisition of cis-regulatory elements in the ethylene biosynthesis genes to complete a positive feedback loop and finally evolve cis-regulatory elements in the downstream ripening genes, which enabled them to be coupled to the loop, could be one of the simplest paths to achieve climacteric fruit ripening. Our findings also suggest that the evolution of climacteric fruit ripening was constrained by the limited set of suitable signalling molecules, genetic or epigenetic building blocks available in the carpel tissues, and WGD provided new resources for plants to circumvent this limit.

**Fig. 5.**
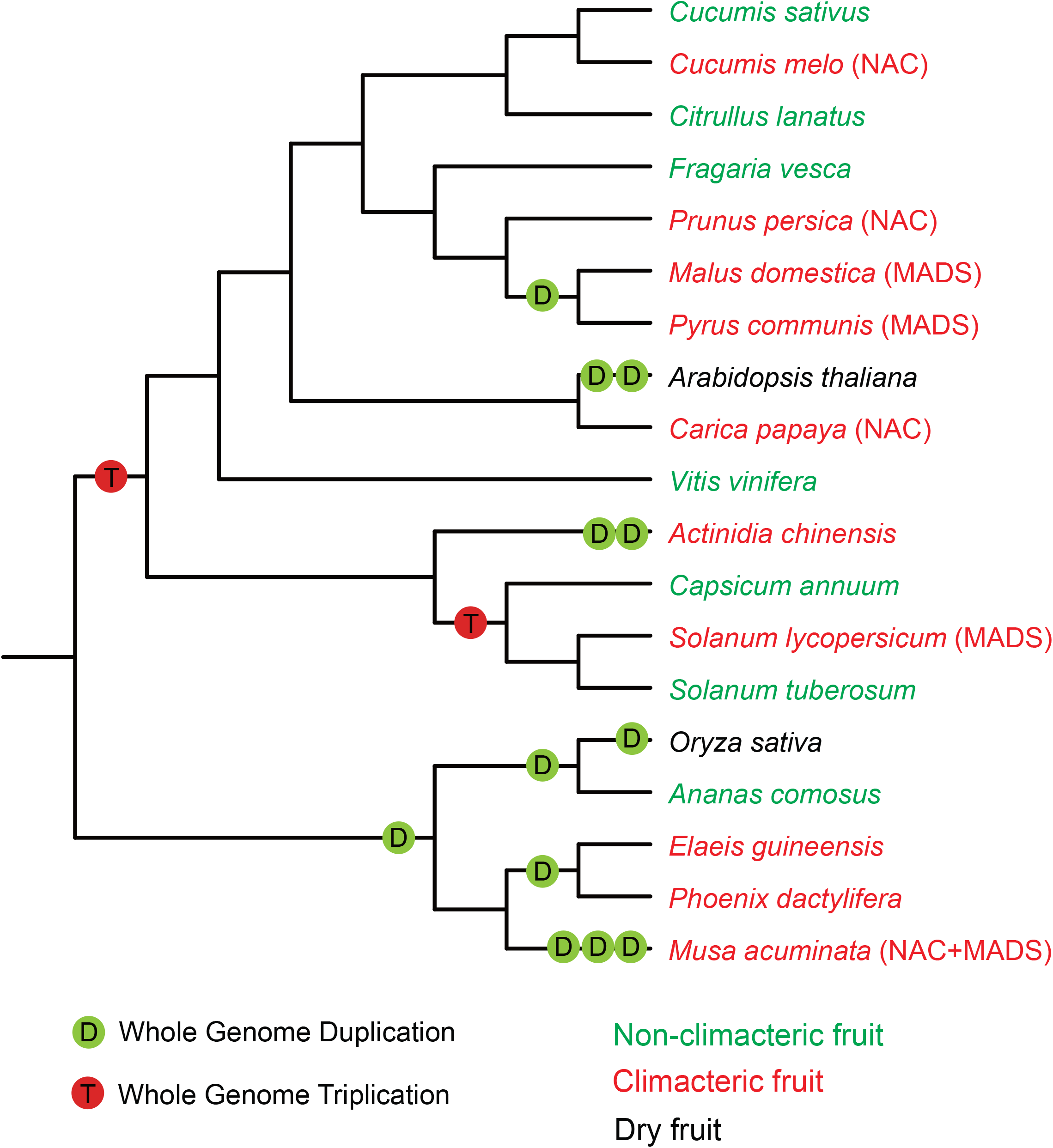
Speciation, fruit ripening types and polyploidization in different angiosperm lineages. Confirmed whole-genome duplications and triplications are shown with circles. Species bearing dry, fleshy climacteric and fleshy non-climacteric fruits are indicated by the color black, red and green, respectively. The three climacteric fruit types are shown in brackets.

The fruitENCODE data also enabled us to systematically compare the epigenetic changes associated ripening in seven climacteric plants. Our data shows widespread promoter DNA methylation and DHS changes, as well as genebody H3K27me3 changes (tables S11 and S15). For example, 17231, 1759 and 630 apple genes are associated with hypo-DMR, tissue-specific DHS and hypo-H3K27me3 in the ripe fruit tissue, respectively. However, these DNA methylation and DHS changes are not always conserved across species. For example, the tomato *CNR* gene promoter is associated with two well-characterized DMRs and tissue-specific DHS (*5*), while they are absent in other species. In addition, for plants that operate a NAC-type positive feedback loop, only the NAC gene in peach is associated with promoter DMR and DHS changes (**Fig. 2**). However, the H3K27me3 changes in the ripening ethylene biosynthesis genes, as well as the ripening NAC and MADS transcription factor genes are highly conserved, and their direct orthologs in *Arabidopsis* and rice also carry similar tissue-specific H3K27me3 marks, suggesting that the ripening epigenetic changes could be originated from existing mechanisms in the ancestral dry fruit species.

Angiosperms became abundant toward the end of the Cretaceous period (~65 Mya). There have been many evolutionary transitions from an ancestral dry fruit to fleshy fruit during this time. It is difficult to trace when these diverse fleshy fruit ripening mechanisms evolved. A common triplication occurred ~71 Mya in the Solanaceae family that includes the climacteric fruit tomato and non-climacteric fruit potato. This suggests that tomato is likely to have acquired its ripening MADS loop after the tomato-potato split. Apple and pear shared a recent WGD event ~40 Mya, and their ripening *MADS, ACS* and *ACO* genes are direct orthologs, which differ from the tomato ones (**figs. S3, S24 and S25**), suggesting they have a common ancestor that bore climacteric fruit. Future study of the ripening regulatory circuits in wild species as well as the relatives of the domesticated crops might shed light on the evolutionary origin of their ripening circuits, and how the non-climacteric fruits make use of their *NAC* and duplicated *MADS* genes.

Banana is a monocot that has diverged from eudicots over 100 Mya. The ripening regulatory system in banana is the most complex, but it is the only monocot fruit with high quality genome that we have examined so far. It has experienced three WGDs, and utilizes a leaf-senescence NAC transcription factor, as well as three MADS-box transcription factors to complete the dual-loop system. However, further study in additional monocot species is required to fully understand how they evolved fleshy fruit ripening.

Ethylene is a stress hormone often involved in tissue senescence and is a gaseous molecule that can easily diffuse. Both attributes make it ideal as the signal controlling fruit ripening in climacteric fruits. However, much less is known about the regulatory mechanisms in nonclimacteric fruits. Study of additional non-climacteric cultivars and mutants and comparison with their climacteric counterparts could provide a new perspective to reassess how nonclimacteric ripening is regulated. Many plant hormones have been implicated in fleshy fruit ripening (*23*). We do observe ripening-associated expression of the ABA biosynthesis genes, particularly in the non-climacteric melon cultivar PI 161375 (**table S23**), which is perhaps not surprising as ABA is also associated with stress and involved in senescence.

The fruitENCODE project has generated a comprehensive dataset for eleven fleshy fruit species, which opens the door for addressing some important problems in agricultural practices. For example, post-harvest loss is a major concern for horticulture produce worldwide, especially in developing countries and is prevalent in modern developed food supply chains as well. However, improvement in shelf life through manipulation of ethylene often leads to reduced quality and nutritional value, which is expected as most of the downstream ripening genes are tightly coupled to the autocatalytic ethylene loop. With a comprehensive annotation of the cis-regulatory elements, and much improved understanding of their ripening regulators, it is now possible design strategies to engineer specific genes’ regulatory element to combat post-harvest losses while retaining or improving quality.

## Acknowledgments

This work is supported by HK UGC GRF-14119814/14104515 and Area of Excellence Scheme AoE/M-403/16 to S.Z., Shenzhen Peacock-KQTD201101 to J.Z., Spanish Ministry of Economy and Competitively grant AGL2015-64625-C2-1-R, Centro de Excelencia Severo Ochoa 2016-2020, and the CERCA Programme/Generalitat de Catalunya to J.G-M, and National Science Foundation IOS-1339287 to Z.F. and J.G. Sequencing data have been deposited in the NCBI Sequence Read Archive under the accession number PRJNA381300.

## Author contributions

S.Z. designed the research, P.L., N.Z., Y.C., B.Z., Y.P., J.F., J.A., N.Y. and J.Z. performed the experiments, S.Y., D.T., S.Z. and J.X. analysed the data, D.G., J.G-M., Z.F., J.G. and S.Z. wrote the paper.

## Competing financial interests statement

The authors declare no conflict of interest.

## References

1 B. G. Coombe, The development of fleshy fruits. Annu. Rev. Plant Physiol. 27, 207–228 (1976).

2 G. B. Seymour, J. E. Taylor, G. A. Tucker, Eds., Biochemistry of Fruit Ripening (Springer Netherlands, Dordrecht, 1993).

3 J. Giovannoni, C. Nguyen, B. Ampofo, S. Zhong, Z. Fei, The epigenome and transcriptional dynamics of fruit ripening. Annu. Rev. Plant Biol. 68, 61–84 (2017).

4 M. T. McManus, The plant hormone ethylene (Wiley-Blackwell, 2012).

5 K. Manning et al., A naturally occurring epigenetic mutation in a gene encoding an SBP-box transcription factor inhibits tomato fruit ripening. Nat. Genet. 38, 948–952 (2006).

6 J. Vrebalov et al., A MADS-box gene necessary for fruit ripening at the tomato ripening-inhibitor (rin) LOCUS. Science 296, 343–346 (2002).

7 S. Zhong et al., Single-base resolution methylomes of tomato fruit development reveal epigenome modifications associated with ripening. Nat. Biotechnol. 31, 154–159 (2013).

8 E. J. Mcmurchie, W. B. Mcglasson, I. L. Eaks, Treatment of fruit with propylene gives information about the biogenesis of ethylene. Nature 237, 235–236 (1972).

9 R. Liu et al., A DEMETER-like DNA demethylase governs tomato fruit ripening. Proc. Natl. Acad. Sci. U.S.A. 112, 10804–10809 (2015).

10 The Tomato Genome Consortium, The tomato genome sequence provides insights into fleshy fruit evolution. Nature 485, 635–641 (2012).

11 J. C. Pech, M. Bouzayen, A. Latché, Climacteric fruit ripening: Ethylene-dependent and independent regulation of ripening pathways in melon fruit. Plant Sci. 175, 114–120 (2008).

12 V. Paul, R. Pandey, G. C. Srivastava, The fading distinctions between classical patterns of ripening in climacteric and non-climacteric fruit and the ubiquity of ethylene—An overview. J. Food Sci. Technol. 49, 1–21 (2012).

13 J. B. Golding, D. Shearer, S. G. Wyllie, W. B. McGlasson, Application of 1-MCP and propylene to identify ethylene-dependent ripening processes in mature banana fruit. Postharvest Biol. Technol. 14, 87–98 (1998).

14 C. Martel, J. Vrebalov, P. Tafelmeyer, J. J. Giovannoni, The tomato MADS-box transcription factor RIPENING INHIBITOR interacts with promoters involved in numerous ripening processes in a COLORLESS NONRIPENING-dependent nanner. Plant Physiol. 157, 1568–1579 (2011).

15 T. Kunieda et al., NAC family proteins NARS1/NAC2 and NARS2/NAM in the outer integument regulate embryogenesis in Arabidopsis. Plant Cell 20, 2631–2642 (2008).

16 Y. Guo, S. Gan, AtNAP, a NAC family transcription factor, has an important role in leaf senescence. Plant J. 46, 601–612 (2006).

17 P. Ríos et al., *ETHQV6.3* is involved in melon climacteric fruit ripening and is encoded by a NAC domain transcription factor. Plant J. 91, 671–683 (2017).

18 R. Pirona et al., Fine mapping and identification of a candidate gene for a major locus controlling maturity date in peach. BMC Plant Biol. 13, 166 (2013).

19 A. D’Hont et al., The banana (*Musa acuminata*) genome and the evolution of monocotyledonous plants. Nature 488, 213–217 (2012).

20 C. Liang et al., OsNAP connects abscisic acid and leaf senescence by fine-tuning abscisic acid biosynthesis and directly targeting senescence-associated genes in rice. Proc. Natl. Acad. Sci. U.S.A. 111, 10013–10018 (2014).

21 S. R. Choudhury, S. Roy, A. Nag, S. K. Singh, D. N. Sengupta, Characterization of an AGAMOUS-like MADS box protein, a Probable constituent of flowering and fruit ripening regulatory system in banana. PLoS One. 7, e44361 (2012).

22 T. Elitzur et al., Banana MaMADS transcription factors are necessary for fruit ripening and molecular tools to promote shelf-life and food security. Plant Physiol. 171, 380–91 (2016).

23 R. Kumar, A. Khurana, A. K. Sharma, Role of plant hormones and their interplay in development and ripening of fleshy fruits. J. Exp. Bot. 65, 4561–4575 (2013).

